# Beyond benchmark accuracy: machine-learning turnover-number predictors require system-level validation

**DOI:** 10.64898/2026.08.28.747816

**Authors:** María José Rimón Martínez, Johann Lottermoser, Alexandre Theodore Charles Bouillon, Wim Vranken, Leopold Zehetner, Jürgen Zanghellini

## Abstract

Enzyme turnover numbers (k_cat_) are essential for kinetic models and enzyme-constrained genome-scale metabolic models (ecGEMs), but measured values are sparse and therefore increasingly estimated using machine learning (ML). Although these predictors are commonly evaluated by global regression metrics, their practical utility depends on how errors propagate through downstream models.

We benchmarked six current k_cat_ predictors on a curated BRENDA-derived dataset and five of them on EnzyExtract. To assess the influence of training-set proximity, we compared each benchmark with the available training data for each predictor. We then used the predicted k_cat_ values to parameterize ecGEMs of *Saccharomyces cerevisiae* and evaluated growth predictions across 19 conditions. We find that benchmark accuracy is moderate even on the BRENDA-derived dataset and drops sharply on EnzyExtract, where all predictors achieve ***R*^2^** values of 0.20 or lower. This decline is accompanied by substantially lower overlap between the benchmark and training datasets, with exact sequence matches ranging from 24 % to 78 % for BRENDA compared with 9 % to 26 % for EnzyExtract. However, that overlap alone does not explain differences in generalization across predictors.

Moreover, downstream performance is also not explained by benchmark ranking. Across 19 conditions, none of the tool-specific ecGEMs consistently reproduces the experimentally observed variation in growth. In glucose minimal medium, the weakest benchmark performer yields the most accurate growth prediction in the downstream ecGEMs, whereas higher-ranked predictors produce larger deviations in growth. We trace this mismatch to localized errors at high-leverage positions in yeast’s metabolic network, where underpredicted mitochondrial ADP/ATP carrier turnover numbers restrict adenine nucleotide exchange and impose an apparent limitation on cytosolic ATP supply. Relaxing this constraint shifts predicted growth toward the experimental reference.

Thus, ML-derived k_cat_ values can affect not only quantitative growth predictions but also the phenotype a mechanistic model appears to identify. These results argue for application-driven validation of biological parameter predictors in the downstream systems they are intended to support.

## 1 Introduction

The catalytic turnover number k_cat_ quantifies the maximum number of substrate molecules converted to product per active site per unit time under saturating substrate conditions. It is one of the central kinetic parameters used to describe enzyme activity and is increasingly important for systems biology [1, 2], synthetic biology [3], and metabolic engineering [4]. In particular, k_cat_ values are required to parameterize kinetic models [5, 6] and enzyme-constrained genome-scale metabolic models (ecGEMs) [7, 8], where they determine how much enzyme is needed to sustain a given metabolic flux.

Despite their importance, experimentally measured k_cat_ values remain sparse [9]. Curated resources such as BRENDA [10, 11] and SABIO-RK [12, 13] contain many kinetic measurements, but the vast majority of enzymes still lack experimentally determined turnover numbers. Moreover, k_cat_ values are condition-dependent and can vary substantially for the same enzyme–substrate pair depending on assay conditions, enzyme source, and data curation choices. This creates a fundamental bottleneck for large-scale kinetic parameterization: k_cat_ values are needed at proteome scale, but they cannot currently be measured at that scale.

Machine learning (ML) approaches have therefore become an attractive route for predicting missing k_cat_ values [14]. Recent tools use enzyme sequences, substrate or reaction representations, protein language-model embeddings, structural information, and multimodal architectures to estimate turnover numbers for enzyme–reaction pairs [15–20]. These models are typically evaluated by comparing predicted and experimental k_cat_ values on benchmark datasets using global regression metrics such as *R*^2^, correlation coefficients, mean absolute error (MAE), or root mean squared error (RMSE). Such benchmarks are useful for assessing isolated prediction accuracy, but they do not necessarily determine whether predicted kinetic parameters are useful in downstream mechanistic models.

This distinction is particularly important for ecGEMs. In contrast to standard genome-scale metabolic models (GEMs), where enzymes are usually implicit annotations, ecGEMs make the cellular proteome an explicit constraint on metabolic flux. In these models, k_cat_ values directly link enzyme abundance to reaction capacity through enzyme-capacity constraints of the form

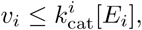

where *v_i_* denotes the flux through reaction *i*, [*E_i_*] the abundance or effective concentration of the catalyzing enzyme, and *k^i^* the corresponding turnover number. Prediction errors in *k^i^* can therefore propagate directly into flux constraints. However, this propagation is not uniform across the network. Errors in inactive or weakly coupled reactions may have little effect, whereas errors in high-flux or growth-limiting reactions can dominate model behavior [21]. Thus, global benchmark accuracy may be poorly aligned with systems-level utility.

Here, we ask whether conventional benchmark rankings identify k_cat_ predictors that are useful in downstream ecGEM simulations. We reproducibly evaluated six current ML tools–CataPro [16], CatPred [15], DLKcat [22], MMKcat [18], TurNuP [19], and UniKP [20]–on a curated BRENDA-derived benchmark and on the largely independent EnzyExtract dataset [23]. We then used tool-specific predictions to parameterize Yeast9-based ecGEMs and assessed their ability to predict growth phenotypes. We find that benchmark rankings fail to predict downstream ecGEM performance, with growth predictions instead shaped by localized kinetic errors at high-leverage network positions.

## 2 Results

### 2.1 Benchmark accuracy remains limited and training-source dependent

We first asked how well current machine-learning models predict enzyme turnover numbers under conventional benchmark settings. We evaluated six executable k_cat_ prediction tools, spanning sequence-, substrate-, reaction-, and structure-aware architectures, on a curated BRENDA-derived benchmark restricted to wild-type entries with all required input features available across tools (Fig. 1a). To account for multiple reported k_cat_ values for the same enzyme-reaction context, duplicate entries were aggregated either by selecting the maximum value or by computing the geometric mean. Both aggregation strategies yielded qualitatively consistent model rankings; therefore, we report results based on the maximum-k_cat_ aggregation strategy unless stated otherwise.

**Fig. 1:**
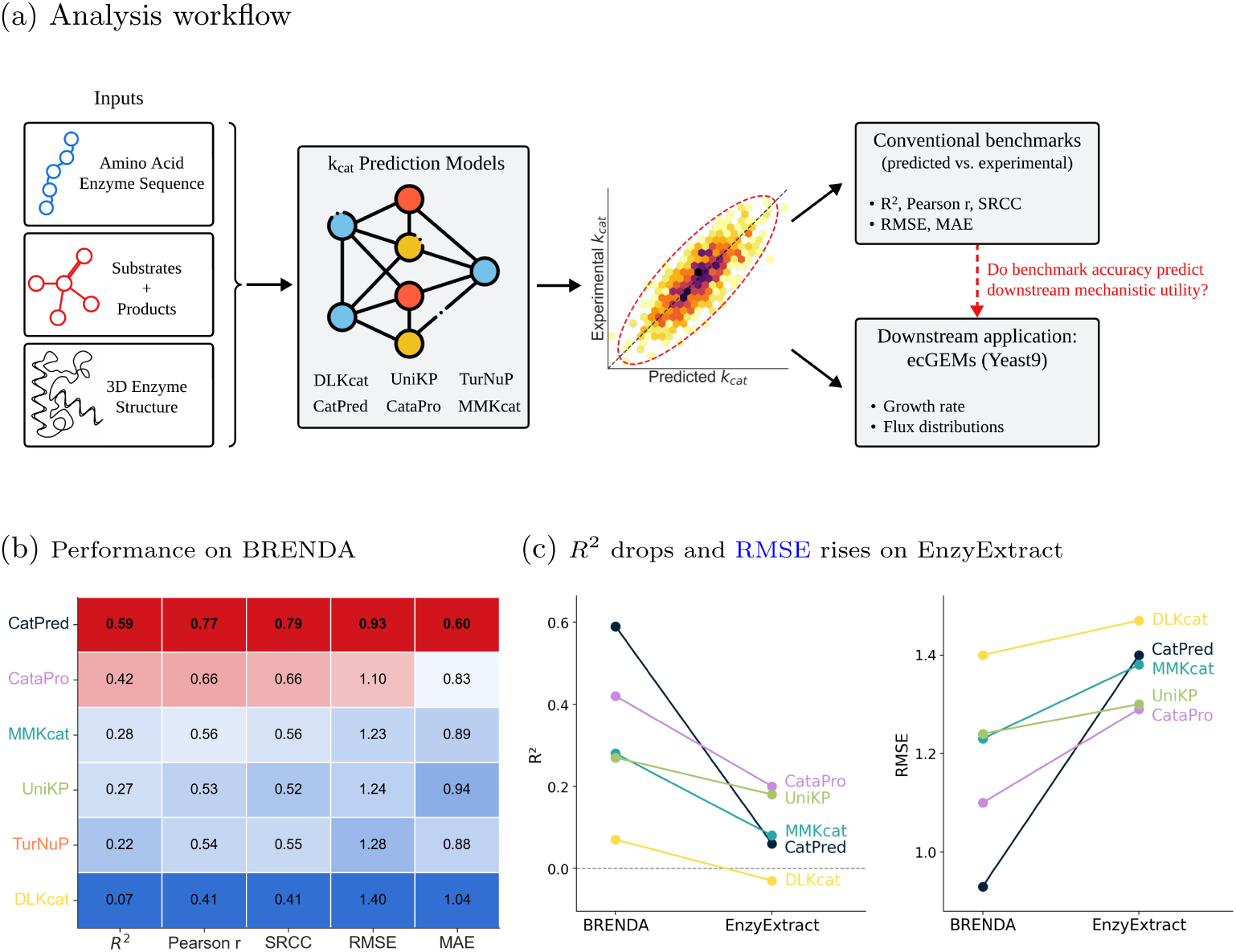
Benchmark performance of machine-learning k_cat_ predictors. (a) Current ML tools typically predict k_cat_ values from enzyme, substrate, reaction, and/or structure information. We compare conventional kinetic benchmark performance with downstream behavior in ecGEMs. (b) Performance on the BRENDA benchmark. Colors indicate relative performance within each metric, with better-performing tools shown in red and worse-performing tools shown in blue. CatPred achieves the highest overall performance but remains limited, reaching *R*^2^ = 0.59. (c) Performance on the EnzyExtract benchmark. TurNuP is omitted because EnzyExtract does not provide the product Simplified Molecular Input Line Entry System (SMILES) required by the model. All other tools show reduced performance on EnzyExtract, with *R*^2^ *≤* 0.20 and higher RMSE values than on the BRENDA-derived benchmark. Detailed predicted-versus-experimental plots and performance statistics for each tool are provided in Supplementary Table SI 2.

Across tools, predictive performance on the BRENDA data set was moderate, and no method achieved uniformly strong agreement with experimental measurements (Fig. 1b). CatPred and CataPro performed best overall, whereas DLKcat showed the weakest performance among the evaluated tools. Even for the best-performing model, however, accuracy remained limited. CatPred reached a Pearson correlation of 0.77 and a Spearman correlation of 0.79, but only an *R*^2^ of 0.59. Prediction errors were substantial, with a MAE of 0.60 and a RMSE of 0.93 on the logarithmic scale, corresponding to typical deviations of approximately 4-fold and 8.5-fold on the original scale. Predicted-versus-experimental scatter plots further revealed a systematic compression of the experimentally observed k_cat_ range, with low experimental values tending to be overestimated and high values underestimated.

To assess performance on a broader, less training-associated dataset (Supplementary Table SI 1), we repeated the evaluation on EnzyExtract [23]. Performance dropped markedly for all evaluated models. *R*^2^ values decreased to 0.20 or below (Fig. 1c), Pearson and Spearman correlations fell below 0.50, and error metrics increased across tools (Supplementary Table SI 2). The strongest decline was observed for CatPred, whose *R*^2^ dropped from 0.59 on the BRENDA-derived benchmark to 0.06 on EnzyExtract. CataPro performed best on EnzyExtract, but still reached only *R*^2^ = 0.20. Thus, conventional benchmark performance is not only limited in absolute terms, but also strongly dependent on the evaluation dataset.

To examine whether the stronger performance on the BRENDA-derived benchmark could reflect greater overlap with the model training data, we compared benchmark sequences with the available training sequences for each predictor. Exact sequence matches are more frequent in BRENDA than in EnzyExtract, ranging from 24 % to 78 % compared with 9 % to 26 % (Supplementary Table SI 1). The largest reductions in exact sequence overlap are observed for CatPred and CataPro, at approximately 50 and 55 percentage points, respectively. For most predictors, the decline in predictive performance broadly tracks the reduction in sequence overlap (Fig. 2). These results suggest that predictive performance for most predictors depends strongly on training-set proximity and deteriorates on less training-associated data. CataPro shows a smaller decline in performance despite its large reduction in sequence overlap and remains the best-performing predictor on EnzyExtract, suggesting comparatively better generalization, although its absolute performance remains limited.

**Fig. 2:**
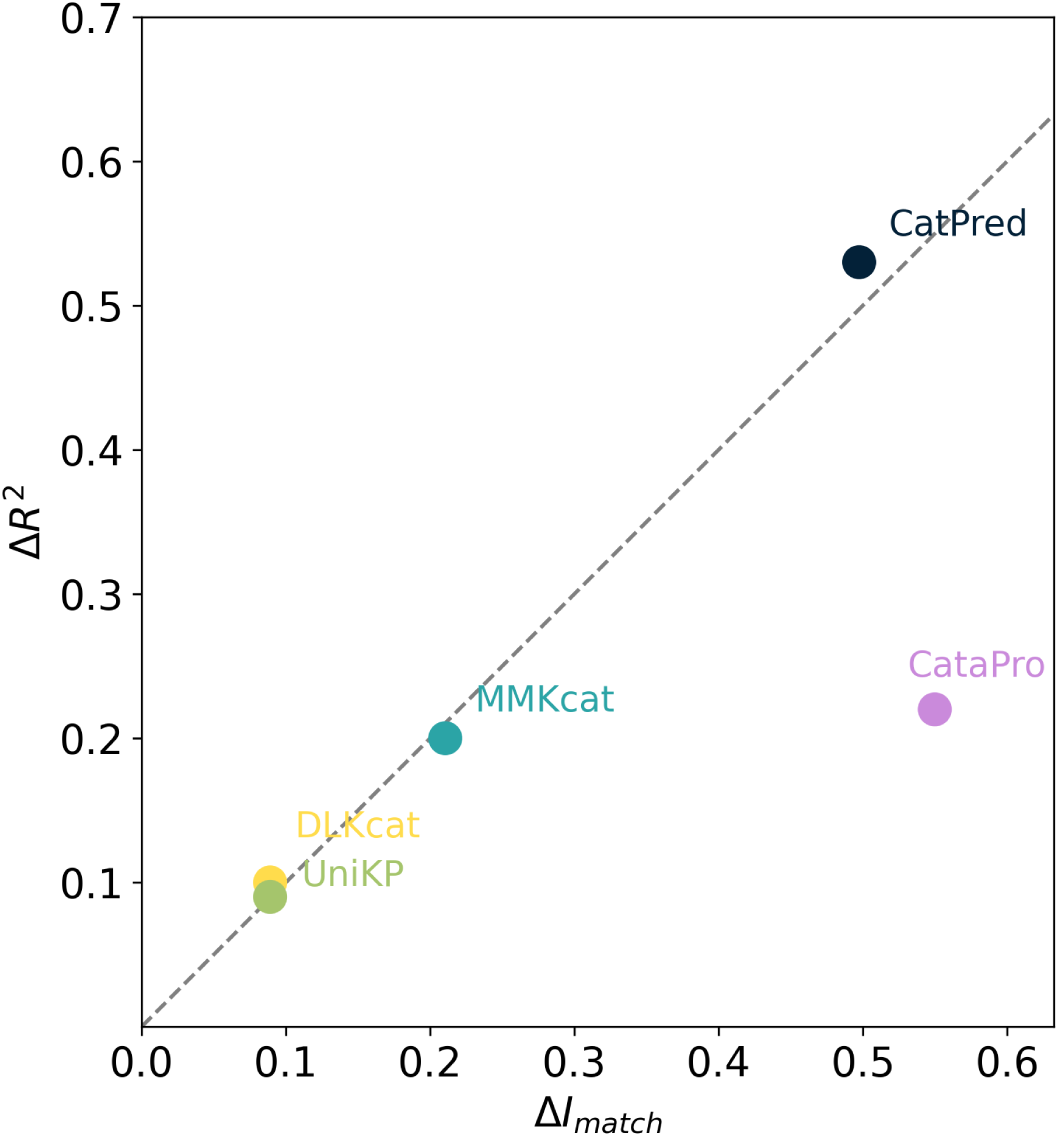
Decline in predictive performance with the reduction in training-set overlap from BRENDA to EnzyExtract. For each predictor, the decrease in *R*^2^ from BRENDA to EnzyExtract, Δ*R*^2^ = 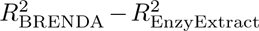, is plotted against the corresponding decrease in the fraction of benchmark sequences with an exact training-set match, Δ*I*_match_ =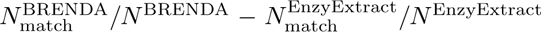. Here, 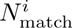 and *N^i^* denote the number of unique benchmark sequences with an exact training-set match and the total number of unique benchmark sequences, respectively, for benchmark *i*. The dashed diagonal marks the 1 : 1 line. Four of the five points lie close to it. However, the line does not represent a theoretical expectation. TurNuP is omitted because EnzyExtract does not provide the product SMILES required by the model.

### 2.2 Tools agree more on failures than successes

We next asked whether tools agree on their most and least accurate k_cat_ predictions. For each tool, predictions were ranked by absolute logarithmic error, and the top and bottom 10 % were selected as the most and least accurate predictions, respectively. Their overlap was then analyzed using set intersection and visualized using UpSet plots (Fig. 3).

**Fig. 3:**
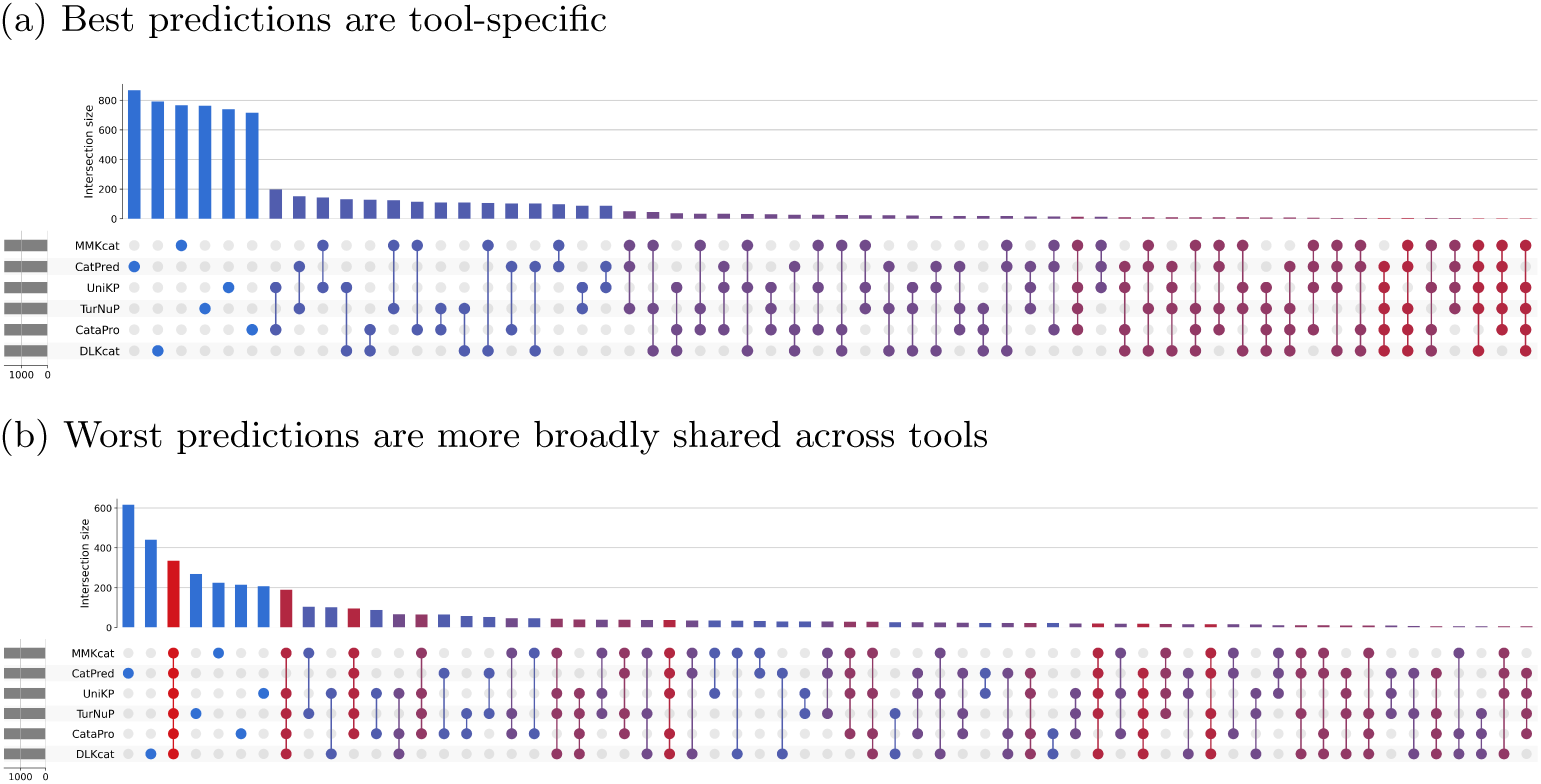
Overlap of best and worst k_cat_ predictions across tools. For each tool, predictions on the BRENDA-derived benchmark were ranked by absolute logarithmic error. The top 10 % most accurate predictions (a) and bottom 10 % least accurate predictions (b) were selected and compared by set-intersection analysis. Intersection-size bars are colored by the number of sets contributing to each intersection, from blue for single-tool intersections to red for intersections shared by all six tools.

For the most accurate predictions, the largest intersections corresponded to single-model subsets (Fig. 3a), while overlaps involving multiple tools were much smaller. In contrast, the least accurate predictions showed stronger agreement: although the two largest subsets were specific to CatPred and DLKcat, the next-largest intersection was shared by all models (Fig. 3b). Thus, high-accuracy predictions were largely tool-specific rather than shared across methods, whereas severe mispredictions included a common subset of difficult-to-predict enzyme–reaction contexts, suggesting shared limitations in the underlying data or molecular representations. However, an overrepresentation analysis of enzyme classes within each intersection set did not reveal any significant enrichment.

### 2.3 ML-parameterized ecGEMs poorly reproduce growth across conditions

Next, we evaluated how k_cat_ predictions propagate to growth phenotypes in a down-stream mechanistic model. Predicted k_cat_ values from each tool were integrated into Yeast9, a GEM of *Saccharomyces cerevisiae*, to construct tool-specific ecGEMs. Maximum growth rates were predicted using flux balance analysis (FBA) across 19 experimental conditions spanning different carbon sources and medium compositions (Fig. 4), using media definitions and corresponding experimental growth rates reported previously [24].

**Fig. 4:**
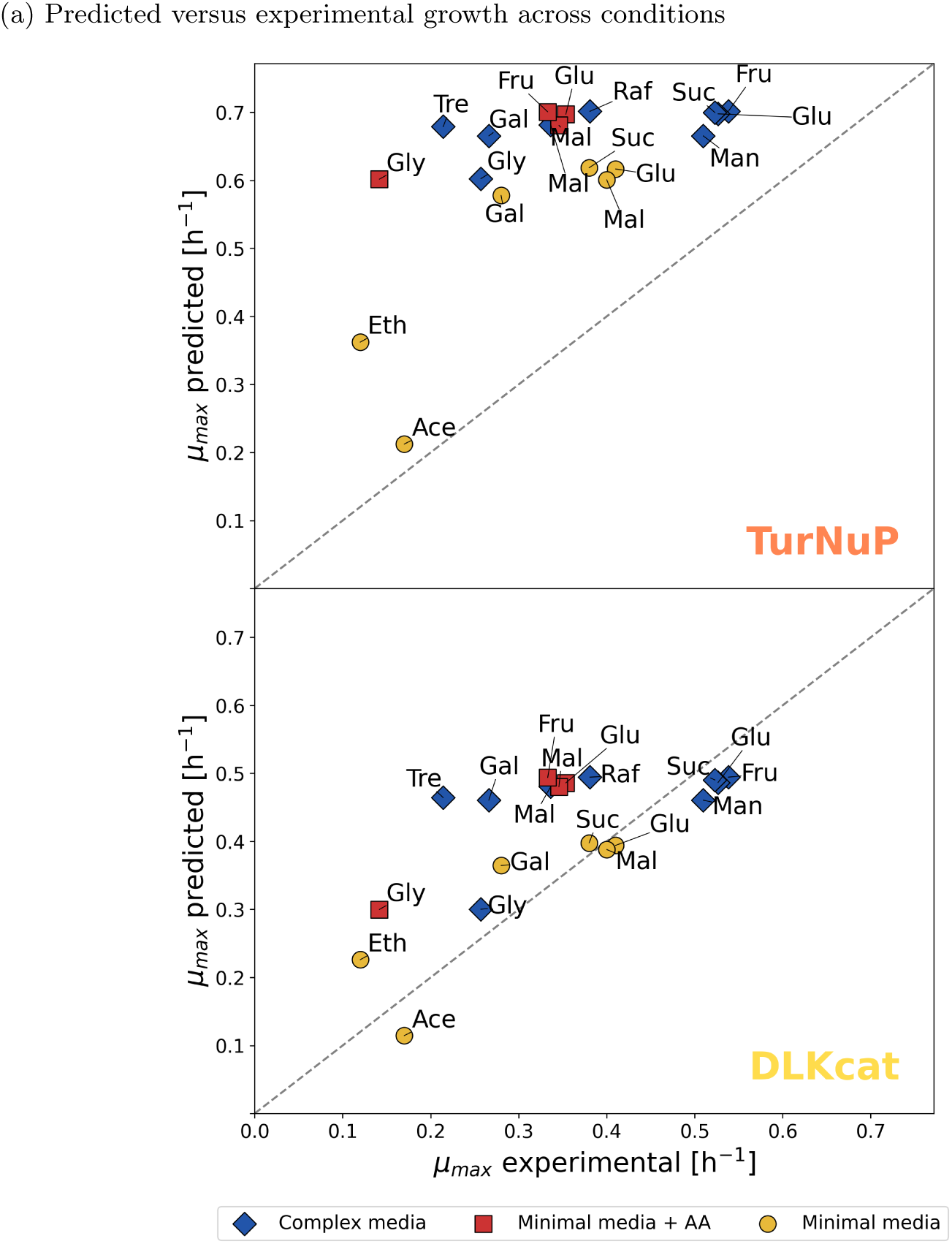
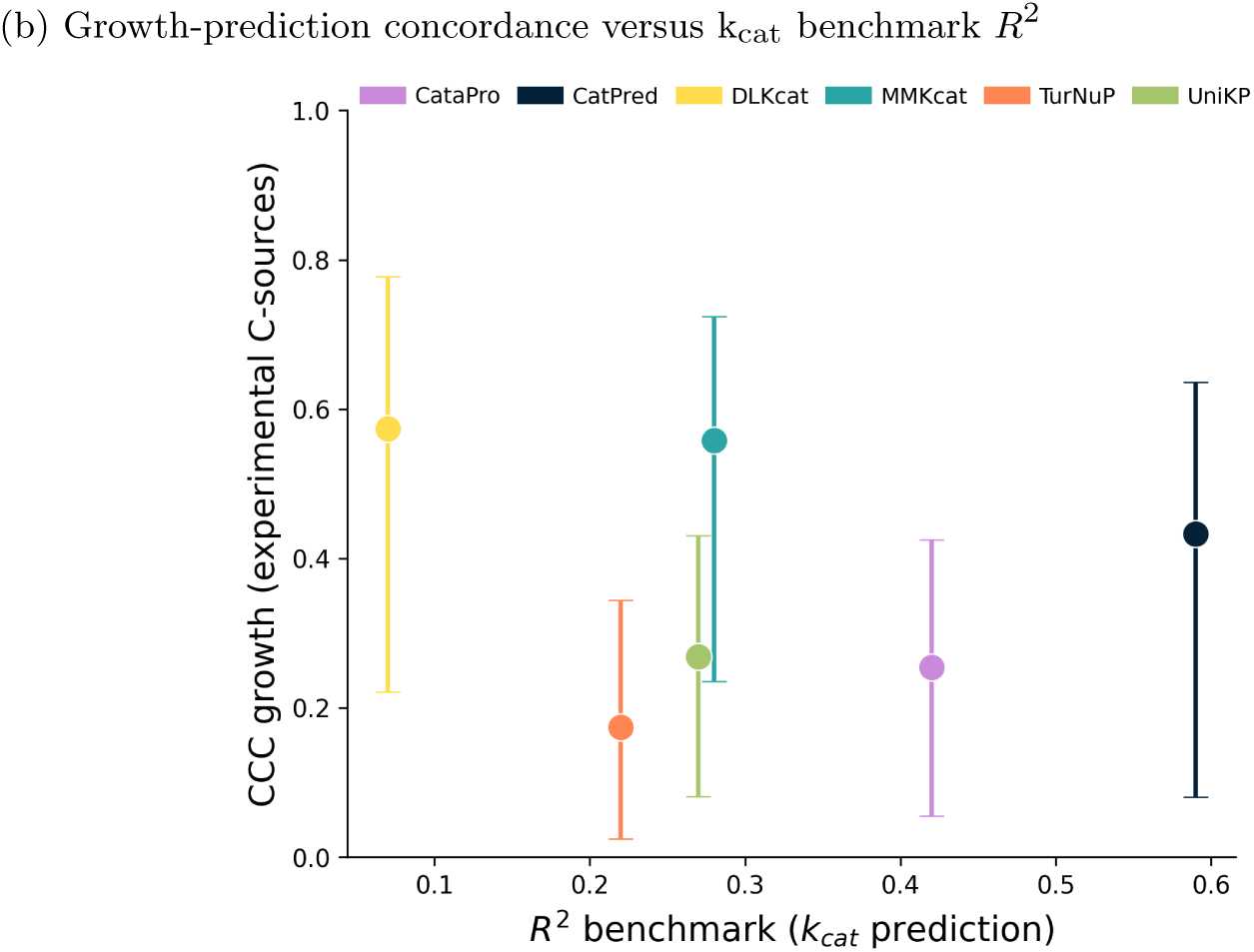
Accuracy of ML-parameterized ecGEMs across multiple conditions. (a) Predicted versus experimental maximum growth rates for DLKcat and TurNuP, representing the highest and lowest across-condition concordance, respectively. Points represent individual conditions; marker color and shape indicate medium composition, and labels denote carbon source. Results for the remaining predictors are shown in Supplementary Fig. SI 1. Fig. 4: Accuracy of ML-parameterized ecGEMs across multiple conditions (continued). (b) Across-condition agreement between predicted and experimental growth rates, quantified using Lin’s concordance correlation coefficient (CCC), versus BRENDA-derived k_cat_ benchmark *R*^2^. Each point represents one prediction tool; vertical error bars indicate 95 % bootstrap intervals obtained from 10 000 resamples of the 19 experimental conditions.

None of the tool-specific ecGEMs consistently reproduce the experimentally observed variation in growth across conditions (Fig. 4a). Across-condition agreement between predicted and experimental growth rates is limited, with Lin’s CCC ranging from approximately 0.17 to 0.57 across tools (Fig. 4b). DLKcat and MMKcat show the highest concordance, whereas TurNuP shows the lowest. In all cases, the 95 % bootstrap intervals are broad and substantially overlapping. However, even the upper bounds of the bootstrap intervals remain below a CCC of 0.8.

We further find that downstream growth concordance does not align with the ranking from the BRENDA-derived k_cat_ benchmark. DLKcat combines the weakest benchmark *R*^2^ with one of the highest downstream CCC values, whereas CatPred achieves the highest benchmark *R*^2^ but only an intermediate downstream CCC value. Thus, higher benchmark accuracy does not correspond to more concordant condition-dependent growth predictions.

Inspection of the individual conditions (Fig. 4a) revealed that the tool-specific ecGEMs often compressed the experimentally observed range of growth rates, over-predicting slow-growth conditions and underpredicting fast-growth conditions. TurNuP showed an additional systematic bias, overpredicting growth across all tested conditions. Thus, accurate predictions for individual conditions did not translate into accurate predictions of growth variation across conditions.

### 2.4 Localized k_cat_ errors generate a glucose-specific growth bottleneck

To investigate how individual k_cat_ predictions contribute to condition-specific growth errors, we used growth in glucose minimal medium as a case study. For comparison, we included an uptake-constrained GEM without enzyme constraints and ecGEMs parameterized with experimentally derived BRENDA k_cat_ values, log-normally sampled random k_cat_ values drawn from the predicted distribution, or a single global mean k_cat_.

Predicted growth varied substantially among the tool-specific ecGEMs (Fig. 5). All tools except TurNuP underestimated growth, with DLKcat giving the closest prediction to the experimental value, deviating by only 6 %. However, the multi-condition analysis showed that this agreement was specific to glucose and did not reflect consistently superior performance across conditions. Among the reference models, the uptake-constrained GEM and the BRENDA-parameterized ecGEM gave similar predictions, around 20 % below the experimental value. The growth rates predicted by the DLKcat- and MMKcat-parameterized ecGEMs were within approximately 15 % of these reference predictions. The random-k_cat_ controls strongly underpredicted growth, whereas the global-mean-k_cat_ ecGEM overpredicted it by more than sixfold.

**Fig. 5:**
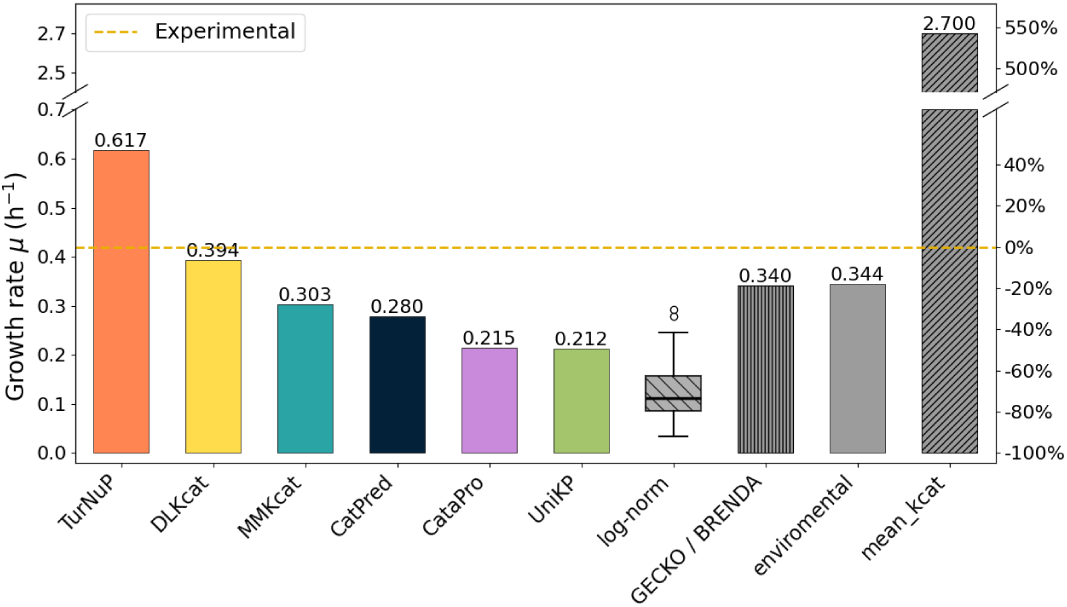
Growth predictions in glucose minimal medium. Maximum growth rates predicted by FBA for tool-specific ecGEMs (colored bars), reference ecGEMs parameterized with log-normally sampled, BRENDA-derived, or global-mean k_cat_ values (patterned bars), and an uptake-constrained GEM without enzyme constraints (gray bar). The dashed line indicates the experimental growth rate of 0.42 h*^−^*^1^ [25].

We asked which enzyme constraints limit growth in the CatPred-parameterized ecGEM. To find out, we repeated FBA while relaxing one enzyme constraint at a time by setting the corresponding k_cat_ value to infinity. Removing most constraints individually has only minor effects on growth (less than 10 %). However, relaxing the constraints associated with the mitochondrial ADP/ATP carrier isoforms Aac1p, Aac2p, and Aac3p (encoded by YMR056C, YBL030C, and YBR085W, respectively) increases the predicted growth rate from 0.28 h*^−^*^1^ to 0.46 h*^−^*^1^ (Fig. 6a). These proteins exchange ATP and ADP across the mitochondrial inner membrane, suggesting that the CatPred-parameterized ecGEM is growth-limited by insufficient cytosolic ATP supply due to restricted mitochondrial adenine nucleotide exchange. Consistent with this bottleneck, DLKcat and TurNuP predict k_cat_ values of approximately 2 *×* 10^5^ h*^−^*^1^ to 5 *×* 10^5^ h*^−^*^1^ for the ADP/ATP carrier isoforms, broadly consistent with available experimental values for Aac2p on the order of 10^5^ h*^−^*^1^ (Fig. 6b). All other tools predict values at least ten-fold lower.

**Fig. 6:**
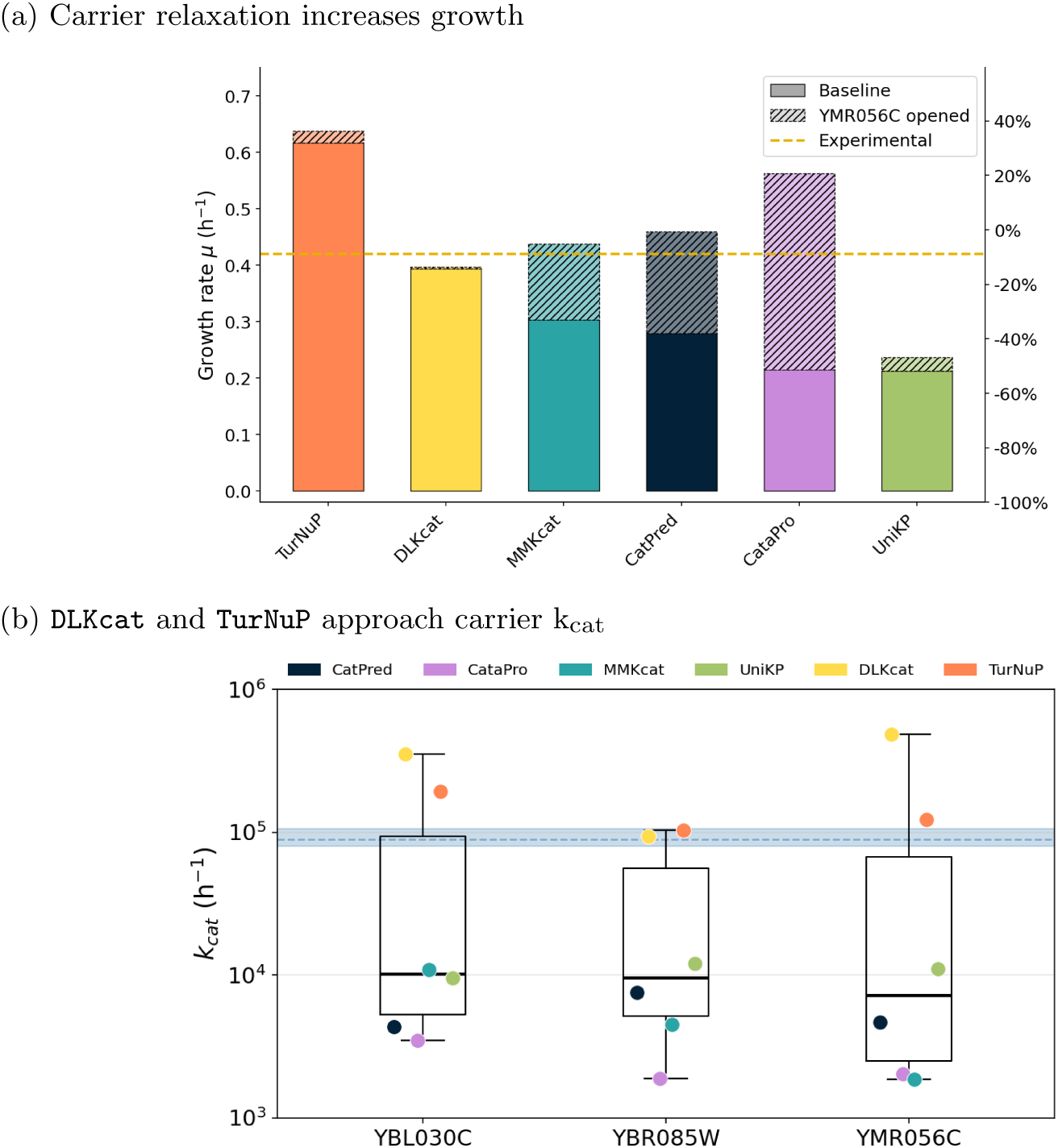
Mitochondrial ADP/ATP carrier constraints strongly contribute to downstream growth deviations. (a) Growth effect of relaxing the YMR056C-associated enzyme constraint across ML-parameterized ecGEMs. Identical growth effects were obtained when the YBL030Cor YBR085W-associated constraints were relaxed instead. Solid bars show original growth rates; stacked hatched bars show the gain after relaxation. The dashed yellow line indicates the experimental reference growth rate of 0.42 h*^−^*^1^ [25]. (b) Predicted k_cat_ values for the mitochondrial ADP/ATP carrier isoforms Aac1p, Aac2p, and Aac3p, encoded by YMR056C, YBL030C, and YBR085W, respectively. The dashed blue line indicates the available experimental k_cat_ value for Aac2p, with uncertainty [26, 27].

This interpretation is supported by the next-largest growth improvement, which occurs when the mitochondrial F1Fo ATP synthase constraint is relaxed. This makes mitochondrial ATP production enzyme-cost-free, freeing enzyme capacity that can instead be allocated to mitochondrial ADP/ATP exchange. Indeed, flux analysis shows increased flux through mitochondrial ADP/ATP exchange, consistent with increased ATP supply to the cytosol. This increases the predicted growth rate to 0.33 h*^−^*^1^, still approximately 22 % below the experimental reference. Together, these results show that downstream growth errors can be driven by localized kinetic bottlenecks that are not captured by global benchmark metrics.

This bottleneck is not unique to the CatPred-parameterized ecGEM. Relaxing the Aac1p/Aac2p/Aac3p-associated constraint substantially increases growth in MMKcat, CatPred, and CataPro, with the largest relative increase observed for CataPro, where predicted growth more than doubles (Fig. 6a). These are the same models for which the corresponding k_cat_ values are predicted at least ten-fold below the available experimental value for Aac2p [26, 27] (Fig. 6b). In contrast, relaxing this constraint has only a small effect in TurNuP and almost no effect in DLKcat, consistent with their higher predicted k_cat_ values for the ADP/ATP carrier isoforms. UniKP, however, shows only a minor growth increase despite low predicted carrier k_cat_ values, indicating that its growth phenotype is dominated by a different limiting constraint.

## 3 Discussion

ML models are increasingly used to predict biological properties and parameters that cannot be measured at scale [28–31]. Their performance is commonly assessed against held-out experimental measurements using global regression metrics [19, 22]. However, performance measured on conventional benchmarks does not necessarily reflect how well a model generalizes to data that differ from its training set.

For example, Kroll and Lercher showed that DLKcat predictions become worse than a constant mean-k_cat_ baseline for enzymes sharing less than 60 % sequence identity with the training set and fail to capture the experimentally observed variation among mutants absent from the training set [32]. This pattern is consistent with our data, as the BRENDA-derived benchmark showed both greater overlap with the predictors’ training data and stronger predictive performance than EnzyExtract (Fig. 2). However, the reduced performance on EnzyExtract should not be attributed exclusively to limited model generalization. Although EnzyExtract was extensively validated against manually curated data and showed strong agreement with matched BRENDA entries [23], residual errors from automated extraction and broader differences in assay and data distributions may also contribute to the observed performance differences.

Likewise, Zheng et al. found poor generalization of current k_cat_ predictors across temporal hold-out data, novel enzymes, deep mutational scans, and enzyme–inhibitor pairs, with predictive ranking deteriorating for sequences increasingly dissimilar from the training data [33]. Alwer and Fleming showed that conclusions about the performance of enzyme kinetic predictors depend strongly on the evaluation strategy, with more stringent sequence-exclusion schemes providing a more challenging assessment of generalization [34]. Similar concerns have been raised for other machine learning applications in biology, where reducing train–test similarity or evaluating models on data generated after model development substantially lowers apparent performance [35, 36].

Our results show that this problem extends beyond the accuracy of the k_cat_ predictions themselves. Even when k_cat_ predictors can be ranked by their agreement with experimental measurements, this ranking alone does not determine their utility once the predictions are used in a downstream mechanistic model.

The key issue is that mechanistic models differ in their sensitivity to individual parameters. Error location can therefore matter more than average error. In constraint-based GEMs, errors in peripheral or inactive reactions may have little effect on the predicted phenotype, whereas errors in high-leverage reactions can dominate model behavior and interpretation [21, 37]. Consequently, two k_cat_ predictors with similar overall errors can have very different downstream consequences if their errors occur at different positions in the metabolic network. Conversely, a predictor with poorer average benchmark accuracy may perform comparatively well for a particular phenotype if its predictions are sufficiently accurate at the constraints that control that phenotype. In this study, underpredicted k_cat_ values for the mitochondrial ADP/ATP carrier isoforms restrict adenine nucleotide exchange and limit cytosolic ATP supply in the CatPred-parameterized ecGEM, suggesting an ATP-supply limitation. The DLKcat-parameterized ecGEM, in contrast, shows little sensitivity to relaxing the same constraint and would therefore not support the same biological interpretation. Thus, predicted k_cat_ values may not only alter quantitative growth predictions but also change which process the mechanistic model identifies as growth-limiting. This does not imply that DLKcat is generally superior to the other predictors. Its comparatively accurate prediction for glucose growth instead illustrates that performance in a specific downstream condition can depend strongly on a small number of influential kinetic constraints. The specific bottleneck may be context-dependent, with other conditions or organisms potentially exposing high-leverage constraints in processes such as redox balancing, biomass precursor synthesis, transport, or other high-flux metabolic reactions. The principle it illustrates is more general. Localized prediction errors at high-leverage constraints can disproportionately shape downstream model behavior and interpretation.

Recent work provides complementary evidence that improved k_cat_ prediction accuracy does not necessarily translate into comparable improvements in downstream enzyme-constrained modeling [34]. Our bottleneck analysis provides a mechanistic example of how such a disconnect can arise.

The example of the mitochondrial ADP/ATP carrier isoforms also cautions against simple consensus strategies. DLKcat and TurNuP are closest to the available experimental estimate for Aac2p but lie far from the center of the cross-tool distribution, so averaging across predictors would not necessarily recover experimentally supported values. More generally, agreement among predictors does not by itself establish reliability. Prediction-specific uncertainty may provide additional information [38], particularly when considered together with downstream model sensitivity. For predictors based on protein language-model embeddings, uncertainty in the underlying protein representation may provide an additional source of uncertainty, as poorly represented proteins have been associated with reduced performance in downstream prediction tasks [39].

All evaluated predictors compress the experimentally observed k_cat_ range to varying degrees, shifting predictions toward central values. Such shrinkage reduces accuracy at the extremes [40]. In a mechanistic model, however, its consequences depend strongly on where these errors occur, because underprediction of high-k_cat_ enzymes at high-leverage reactions can introduce restrictive kinetic bottlenecks (Fig. 6).

Application-driven benchmarks should therefore complement conventional regression benchmarks. This conclusion is supported by recent work in enzyme engineering, where kinetic-parameter predictors that performed well on database-derived benchmarks lost much of their advantage across independent mutagenesis datasets, with prospective experimental validation further revealing differences between benchmark performance and practical utility [41]. For enzyme kinetic prediction, evaluation should therefore address several complementary aspects. Predictors should be tested on kinetic datasets that are as independent as possible from their dominant training sources, and their errors should be examined beyond aggregate regression statistics, including systematic biases, severe underpredictions, and disagreement between methods. At the same time, experimentally measured k_cat_ values are generally obtained in vitro and may not directly correspond to effective catalytic capacities in vivo [9, 42]. Agreement with experimental k_cat_ values should therefore not be regarded as the sole criterion for assessing the usefulness of a predictor in cellular models. Complementary approaches that optimize kinetic parameterization directly within enzyme-constrained models [43] further illustrate the value of evaluating kinetic parameters in their systems-level context.

When predicted parameters are intended for use in mechanistic models, evaluation should additionally extend to the resulting system-level behavior. In this setting, prediction errors could be interpreted in the context of model sensitivity because an error in an inactive or weakly constrained reaction may have little consequence, whereas a similar error in a growth-limiting reaction can dominate the predicted phenotype. Conventional error metrics could therefore be complemented by application-dependent measures that place greater weight on parameters associated with active reactions that carry high flux, or strongly influence the phenotype of interest. More generally, combining parameter uncertainty with model sensitivity may help identify predictions for which improved measurements or more accurate estimates would have the greatest impact on model reliability [44, 45].

Extending such analyses across model formulations, organisms, environmental conditions, and phenotypes will be important for determining whether the high-leverage constraints identified by system-level validation are context-specific or reveal recurring vulnerabilities of kinetic parameterization.

## 4 Materials and methods

### 4.1 Selection and implementation of k_cat_ prediction tools

We surveyed eight published tools for predicting k_cat_ and related kinetic parameters (Table 1). The tools differ in their predicted quantities, input requirements, training-data provenance, public availability, fine-tuning support, and implementation requirements. To ensure reproducible benchmarking, inclusion required that a tool could be installed and executed using the publicly available materials. Six tools met this criterion; NNKcat was excluded because files required for model execution were missing, whereas DEKP was excluded because its trained prediction models were unavailable.

**Table 1:**
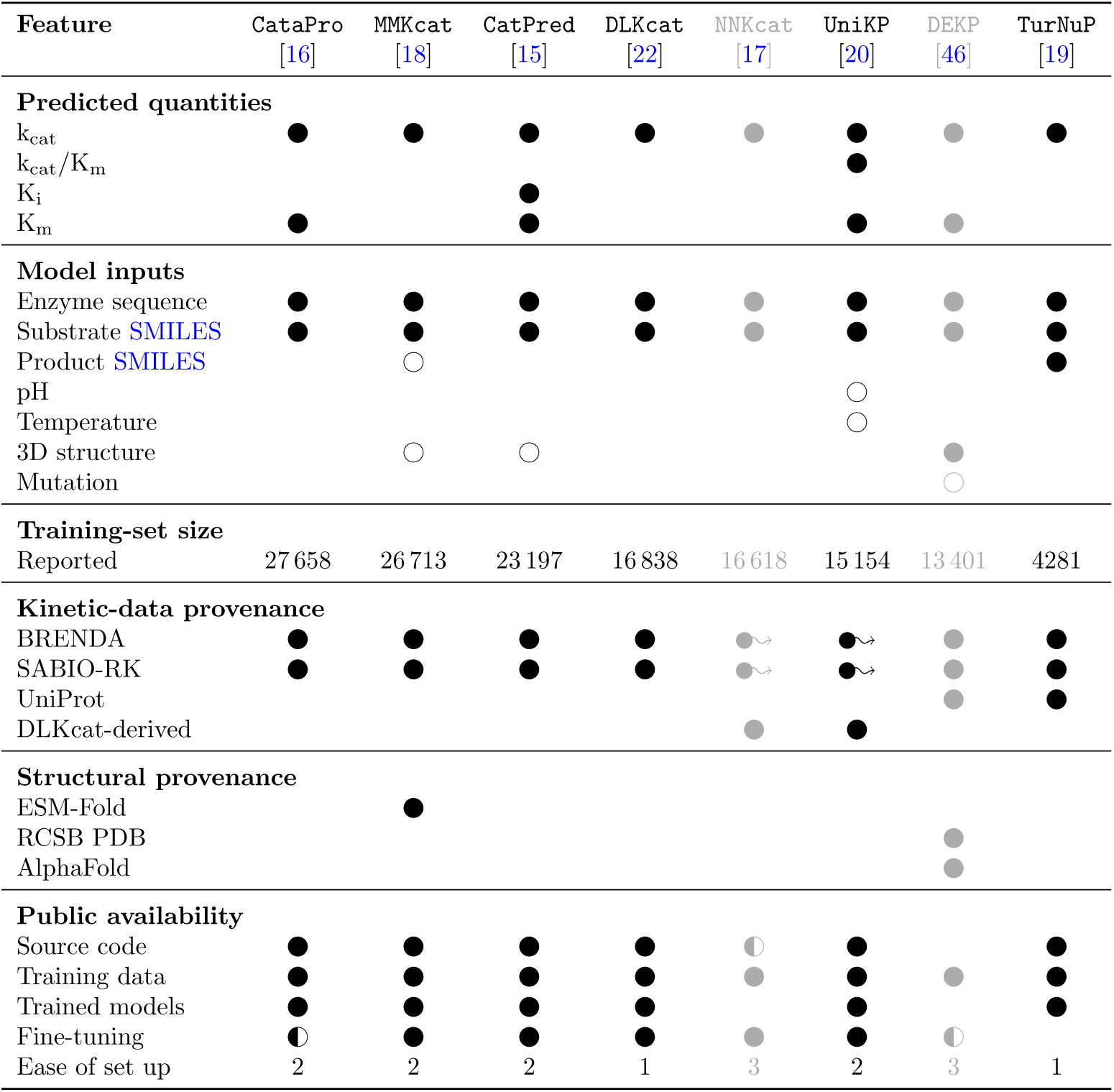
Comparative overview of published k_cat_ prediction tools considered for benchmarking. 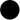 indicates that a feature is available or applicable; 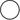 indicates an optional input; 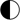 indicates partial availability or support; and 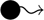∼ indicates provenance inherited indirectly through the DLKcat training set. “Ease of set up” reflects installation and execution in the standardized environment used in this study: 1, setup completed with only minor adjustments; 2, setup required substantial troubleshooting or modification; and 3, setup could not be completed using the publicly available materials. Tools assigned a setup-effort score of 3 were therefore excluded from the benchmark and are indicated by gray text.

Each of the six selected tools was installed in a separate Anaconda environment following the instructions provided by its authors. Additional dependencies and model resources were retrieved and configured where required.

To standardize their use, we developed a unified toolbox that automates environment setup and provides a common interface for k_cat_ prediction across the models. The toolbox also implements the data-generation and curation workflows used to construct benchmark datasets from BRENDA [10] and EnzyExtract [23]. Exported environments, installation scripts, and the toolbox are available at https://github.com/jlott/kcat-prediction-benchmarking.

For benchmarking, all tested models received the enzyme amino acid sequence and substrate SMILES as common required inputs. TurNuP additionally required product SMILES (Table 1). Optional inputs, including pH, temperature, product SMILES for MMKcat, and 3D structural information, were omitted to ensure a consistent input basis. Moreover, previous evaluations found that including 3D structural information did not improve predictive performance [15].

### 4.2 Benchmarking tools with databases

To benchmark the implemented k_cat_ prediction models, multiple data sources were used. The primary benchmark source was BRENDA [10]. It is manually curated from primary literature and has information on enzymes related to 6941 different EC numbers. The second data source that was used is EnzyExtract [23], which is a dataset that was automatically assembled using a custom large language model (LLM) on a large set of scientific publications. EnzyExtract is aimed at ML use cases with each entry containing an experimentally measured k_cat_ value, a substrate, the enzyme sequence and additional metadata.

#### BRENDA

The BRENDA dataset was curated using version“2026 1” of the flatfile. First, each k_cat_ value had to be linked to its respective catalyzing enzyme and the reaction equation. Only enzymes with an associated UniProtId were taken to facilitate a clear amino acid sequence association. The UniProt [47] API service was used to retrieve all relevant sequences. Furthermore, only wild-type entries were retained, as the majority of k_cat_ prediction models were exclusively trained on that type of data. The main BRENDA enzyme database only represents reaction substrates and products using the name found in the original publication. To map between names and SMILES strings which are needed for the input of the k_cat_ prediction models, a multi-step approach was implemented. First, the BRENDA ligand info dataset was downloaded, which contains ligand names and, when available, corresponding InChi strings and ChEBI [48] identifiers. All names found in reaction equations are mapped to the ligand info dataset when possible. Then, ChEBI identifiers are used to directly obtain an accurate SMILES string from the ChEBI API service, using the bioservices python package [49]. Any unresolved molecules remaining after this step are then fed into the MoleculeResolver tool [50], which queries multiple sources for SMILES such as PubChem [51], checks for chemical plausibility and selects a result based on frequency across different sources. All reactions in which the substrate SMILES associated with the k_cat_ value could not be retrieved were discarded. To standardize obtained SMILES they were transformed into their canonical form using the RDKit [52] python package.

To ensure only one k_cat_ per unique reaction, the dataset was deduplicated using a unique combination of enzyme sequence and all substrates. For duplicate reactions, the k_cat_ value was aggregated using either the maximum or the geometric mean. This was done to account for differences in aggregation strategies of the training data across the various k_cat_ prediction models.

#### EnzyExtract

The EnzyExtract database was created with the aim to have a k_cat_data source specifically designed for ML applications with vast amounts of entries. It features one SMILES substrate, the enzyme sequence and the associated k_cat_ value for each entry. The database was downloaded and cleaned for only wild type reactions based on an included precomputed flag. Then it was deduplicated using the same strategies as for the BRENDA dataset.

#### Training-set sequence overlap

To quantify sequence-level overlap between each benchmark and the available training data for each predictor, the benchmark and corresponding training dataset were each reduced to unique enzyme amino-acid sequences. For each unique benchmark sequence *i*, global pairwise alignments were performed against all unique training sequences *j* using Biopython [53], and the highest sequence identity was recorded as the best-match identity,

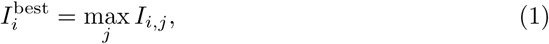

where *I_i,j_* denotes the sequence identity between benchmark sequence *i* and training sequence *j*. Exact sequence matches were defined as benchmark sequences with *I*^best^ = 100 %. The distribution of best-match identities was summarized in Supplementary Table SI 1 in mutually exclusive identity bins of [0, 50)%, [50, 70)%, [70, 90)%, [90, 100)%, and 100 %.

The mean best-match identity was then calculated as

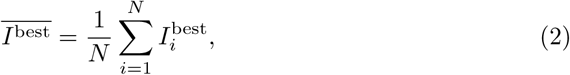

where *N* is the number of unique benchmark sequences.

### 4.3 Construction and analysis of ecGEMs

All constraint-based analyses were performed using COBRA-k (v0.0.8) [5].

#### Base model

Yeast9 [54] was used as the base GEM for *S. cerevisiae*. During preprocessing, COBRA-k split reversible reactions into separate forward and reverse reactions and split reactions associated with isoenzymes (i.e., reactions whose gene–protein– reaction rules contained at least one OR relationship) into separate reaction branches. Each resulting irreversible reaction was associated with at most one branch of a gene– protein–reaction rule. The processed model contained 7122 reactions, of which 4881 were gene-associated.

#### Molecular representation

Yeast9 metabolites are annotated with identifiers from multiple chemical databases. MetaNetX [55] and ChEBI [48] identifiers were selected for SMILES retrieval because they provided the highest coverage. MetaNetX release 4.0 was downloaded on April 8, 2026, and ChEBI was accessed via application programming interface (API) on April 8, 2026. As all MetaNetX metabolites were also represented in ChEBI, MetaNetX was prioritized for overlapping metabolites, while ChEBI was used only for metabolites not covered by MetaNetX. After discarding structures that could not be parsed by RDKit version 2025.9.6 [52], 2068 unique SMILES strings were obtained, enabling reconstruction of substrate and product representations for 2822 reactions.

#### Construction of tool-specific ecGEMs

For the 2822 reactions with complete molecular representations, we extracted all associated protein identifiers and obtained the corresponding amino acid sequences from the NCBI RefSeq annotation of the *S. cerevisiae* S288C genome (assembly accession GCF 000146045.2 SGD annotation version R64–5–1), downloaded in November 2025. After preprocessing, each gene-associated reaction was associated with either an individual enzyme or a multisubunit enzyme complex. For individual enzymes, each k_cat_ predictor was applied using the reaction and sequence inputs described for the benchmark analysis. For multisubunit complexes, k_cat_ was predicted separately for each subunit, and the highest predicted value was retained, resulting in the least restrictive enzyme-capacity constraint.

All tools generated valid k_cat_ predictions for the same set of 2822 reactions with the exception of DLKcat, which could only predict k_cat_ values for 2793 reactions due to its inability to process certain substrate SMILES representations, including wildcard atoms and copper ions. Reactions without a gene association or gene-associated reactions without complete molecular inputs or a valid tool prediction were left unconstrained. Coverage of gene-associated and flux-carrying reactions for each tool-specific ecGEM is reported in Supplementary Table SI 3. Enzyme molecular weights were calculated using Biopython v1.87. For multisubunit complexes, the molecular weights of the constituent subunits are summed internally by COBRA-k to obtain the total molecular weight of the complex. Molecular weights and tool-specific k_cat_ dictionaries were integrated into Yeast9 using the COBRA-k function add thermokinetic data to cobrak model.

#### Construction of reference ecGEMs

In addition to the six tool-specific ecGEMs, we constructed 100 ecGEMs parameterized with independently sampled k_cat_ sets drawn from a log-normal distribution. On their original scale, the pooled predictions from all six tools spanned 4 *×* 10*^−^*^4^ s*^−^*^1^ to 4.38 *×* 10^7^ s*^−^*^1^. The base-10 logarithms of these values were used to estimate a mean of 0.656 and a standard deviation of 0.7. For each replicate, one log-transformed k_cat_ value was sampled independently for each of the 2822 enzyme-constrained reactions using np.random.normal from NumPy v1.21.6 [56]. Finally, the sampled values were converted to the original scale using 10*^x^*.

We additionally constructed one ecGEM in which all enzyme-constrained reactions were assigned the arithmetic mean of the pooled predicted k_cat_ values, 132.4 s*^−^*^1^, and another parameterized with experimentally measured k_cat_ values compiled from BRENDA and provided in the GECKO dataset. For *S.cerevisiae*, the maximum k_cat_ was retained per EC number and mapped to Yeast9 reactions based on their annotated EC numbers; when multiple EC numbers matched a reaction, the maximum corresponding k_cat_ was assigned.

All k_cat_ values were converted to h*^−^*^1^ before integration into Yeast9. All ecGEMs used the COBRA-k default maximum total enzyme-pool constraint of 0.25 g enzyme per g cell dry weight. Reactions without a gene association or without an assigned k_cat_ value were left unconstrained.

Medium compositions were adopted without modification from the original GECKO study [24]. Uptake reactions for metabolites present in the corresponding medium were enabled without an additional finite exchange-flux bound, whereas uptake reactions for metabolites absent from the medium were disabled. All secretion reactions were enabled without additional exchange-flux bounds. Flux through any of these reactions could nevertheless be restricted by enzyme-capacity constraints when an associated k_cat_ value was available.

#### Construction of an uptake-constrained reference GEM

For glucose minimal medium, an uptake-constrained GEM without enzyme constraints was additionally included as a reference. This model used the same medium definition as the ecGEMs, but glucose uptake was limited to 3.9 mmol g*^−^*^1^ h*^−^*^1^ [57]. Uptake of all other metabolites present in the medium and all secretion reactions remained enabled without additional finite bounds.

#### Growth and flux simulations

Maximum growth rates were calculated using FBA with the Yeast9 biomass reaction r 2111 as the objective and the COBRA-k function perform lp optimization. Optimization was performed using glpk solver version 5.0. Flux distributions were obtained using parsimonious flux balance analysis (pFBA) with growth constrained to its maximum value (relaxed by a factor of 10*^−^*^6^ to ensure numerical stability).

To identify growth-limiting enzyme-capacity constraints, each enzyme-specific constraint was relaxed individually by setting its corresponding k_cat_ value to infinity, after which the model was reoptimized. The effect of each relaxation was quantified as

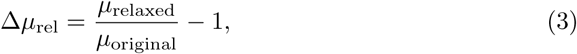

where *µ*_original_ and *µ*_relaxed_ denote the maximum growth rates before and after constraint relaxation, respectively.

#### Statistical evaluation

Across-condition predictive performance was evaluated using Lin’s CCC

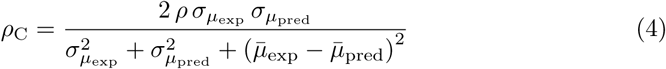

where *µ*_exp*,i*_ and *µ*_pred*,i*_ are the experimental and predicted growth rates for condition *i*, respectively. *ρ* is the Pearson correlation coefficient between *µ*_exp_ and *µ*_pred_ across the 19 experimental conditions; *̄μ*_exp_ and *̄μ*_pred_ are their respective means; and *σ_µ_*_exp_ and *σ_µ_*_pred_ are their respective standard deviations.

To assess the robustness of growth-prediction performance across experimental conditions, we estimated uncertainty in Lin’s concordance correlation coefficient using non-parametric bootstrapping. For each predictor, the 19 experimental conditions were sampled with replacement while preserving the pairing between predicted and experimental growth rates. Lin’s concordance correlation coefficient was recalculated for each of 10 000 bootstrap samples, and the resulting distributions were summarized using the 2.5th and 97.5th percentiles as 95 % bootstrap intervals.

## Supplementary information

Three supplementary tables and one supplementary figure are available online.

## Acknowledgments

Not applicable.

## Ethics approval and consent to participate

Not applicable.

## Consent for publication

We confirm that the manuscript has been read and approved by all named authors and that there are no other persons who satisfied the criteria for authorship but are not listed. We further confirm that the order of authors listed in the manuscript has been approved by all of us.

For open access purposes, the authors have applied a CC BY public copyright license to any author-accepted manuscript version arising from this submission.

## Data availability

Benchmark datasets were derived from publicly available resources, including BRENDA and EnzyExtract, as described in the Methods.

## Materials availability

Not applicable

## Code availability

Code and data-processing workflows used for k_cat_ predictor benchmarking are available at https://github.com/jlott/ kcat-prediction-benchmarking [58].

Code for ecGEM integration and constraint-based modeling analyses is available at https://github.com/mjrimon/ecGEMs analyses [59].

## Author contribution

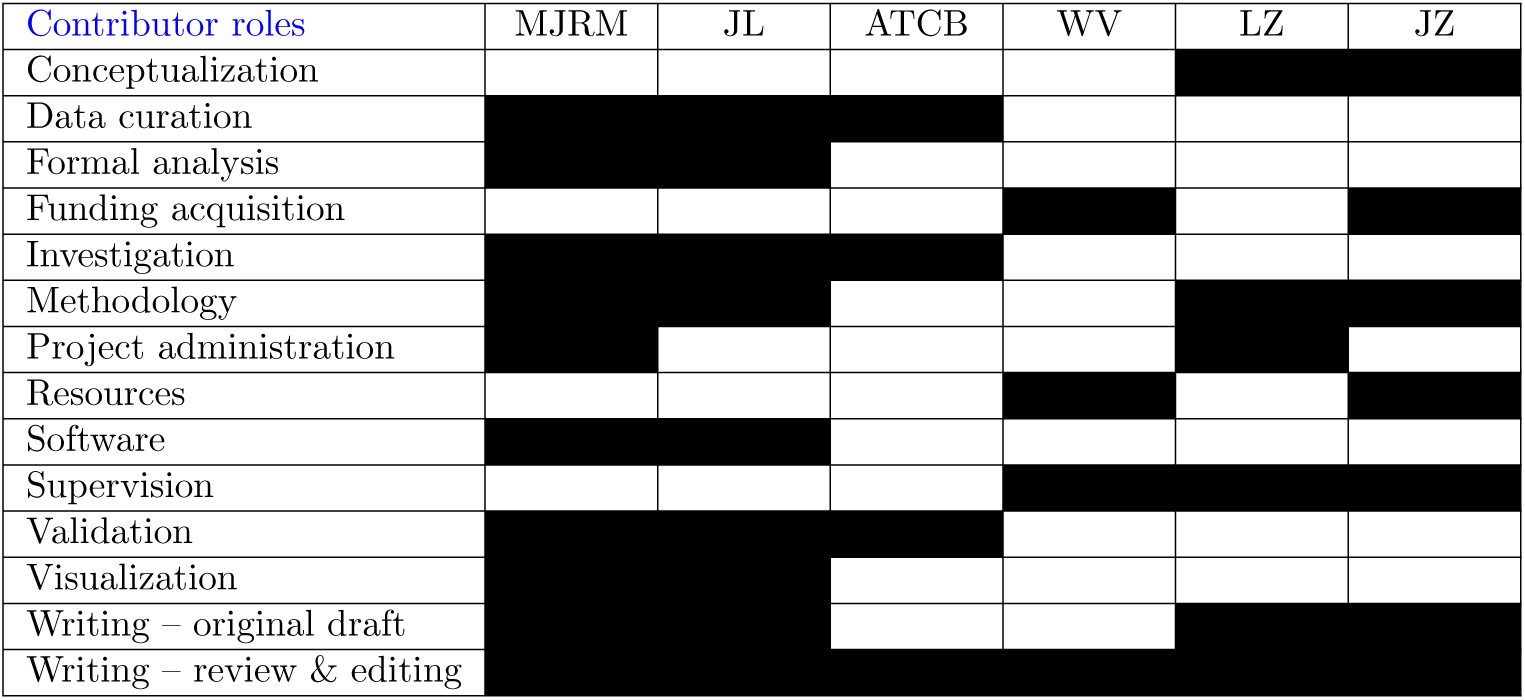

## Supporting information

**Fig. SI 1:**
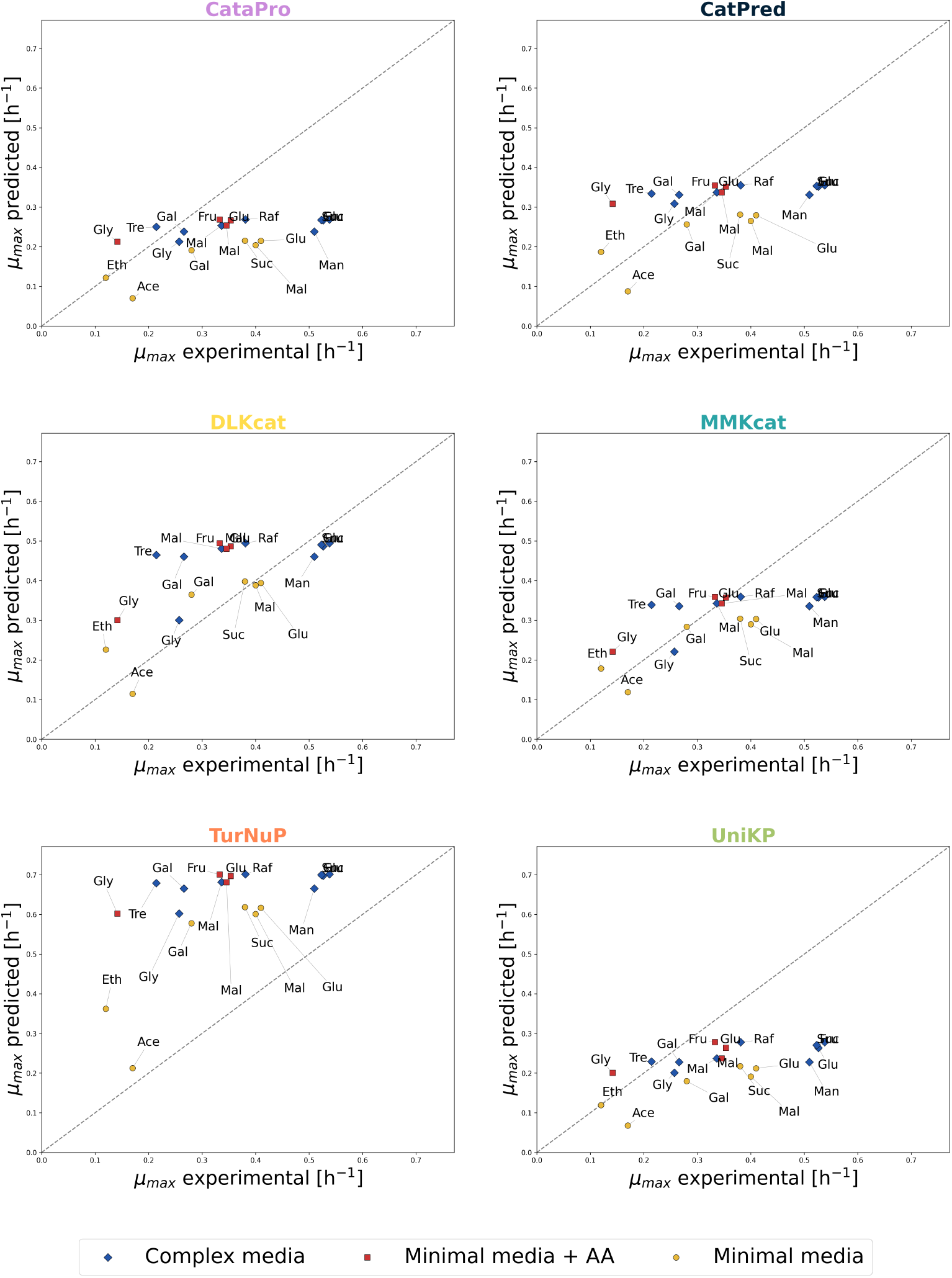
Accuracy of ML-parameterized ecGEMs across multiple conditions. Predicted versus experimental maximum growth rates across carbon sources and medium compositions. Points represent individual conditions; marker color and shape indicate medium composition, and labels denote carbon source.

**Table SI 1:**
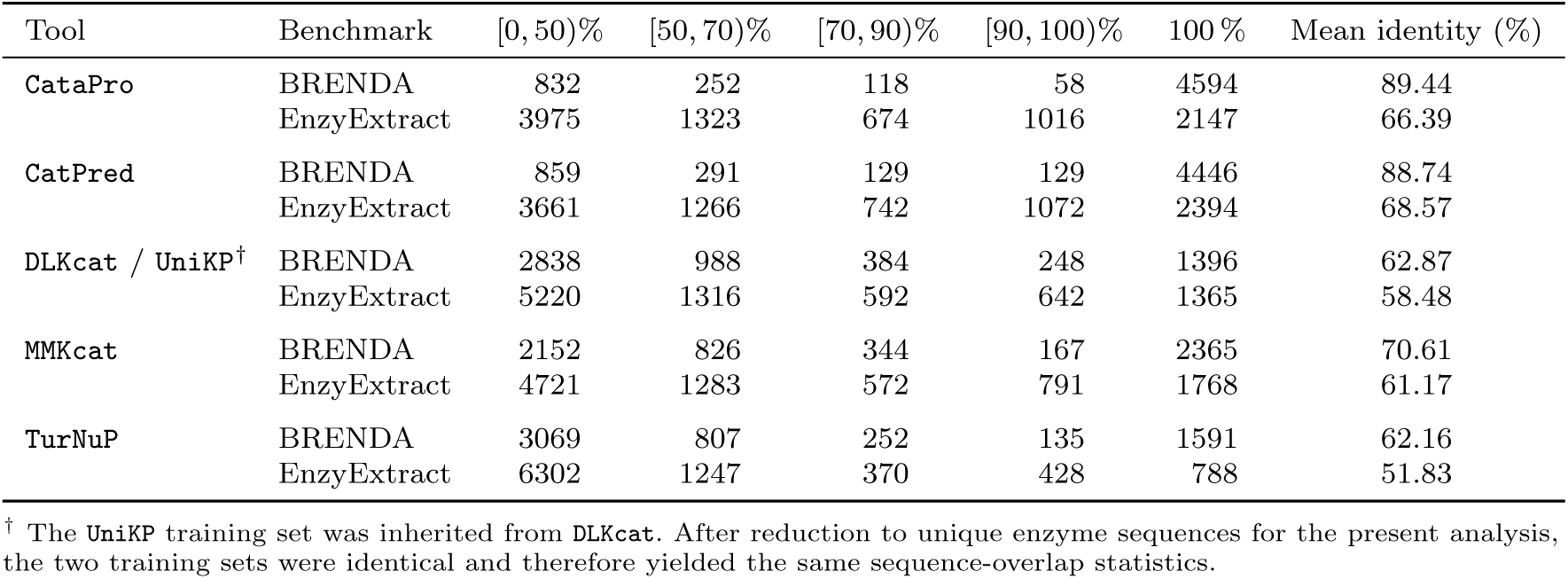
Training-set sequence similarity of the BRENDA and EnzyExtract benchmarks. For each unique benchmark sequence, the closest sequence in the corresponding predictor training set was identified by global pairwise alignment. Values give the number of benchmark sequences within mutually exclusive best-match identity bins. Bin ranges are given in mathematical interval notation, with e.g. [50, 70)% indicating 50 % *≤ x <* 70 %. Exact matches correspond to a best-match identity of 100 %. Mean identity gives the mean best-match sequence identity across all unique benchmark sequences (see Section 4 Materials and methods). In total, the BRENDA and EnzyExtract benchmarks contain 5854 and 9135 unique sequences, respectively. The two benchmark datasets themselves share 1892 exact sequences, corresponding to 20.71 % of unique EnzyExtract sequences and 32.32 % of unique BRENDA sequences.

**Table SI 2:**
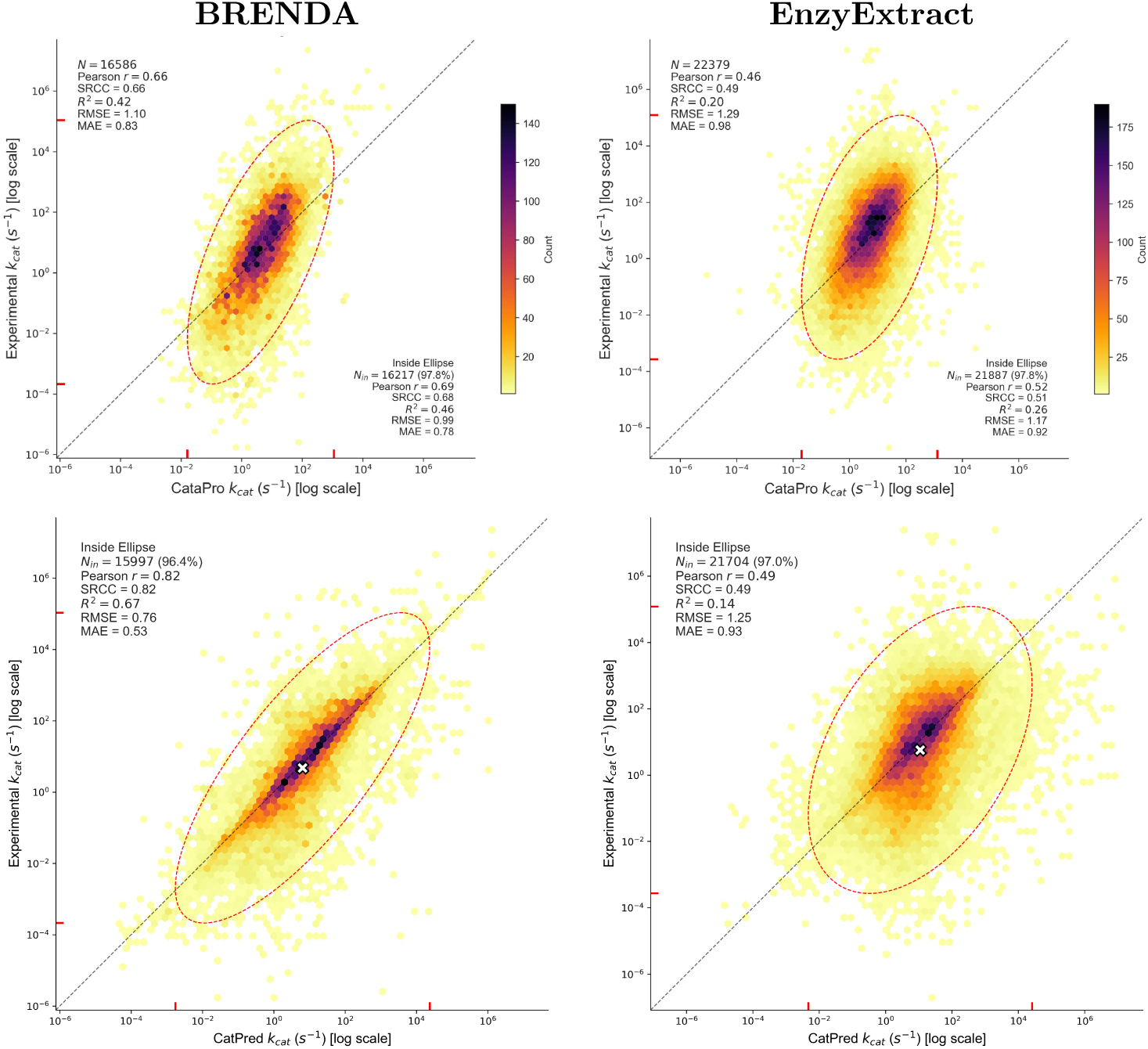
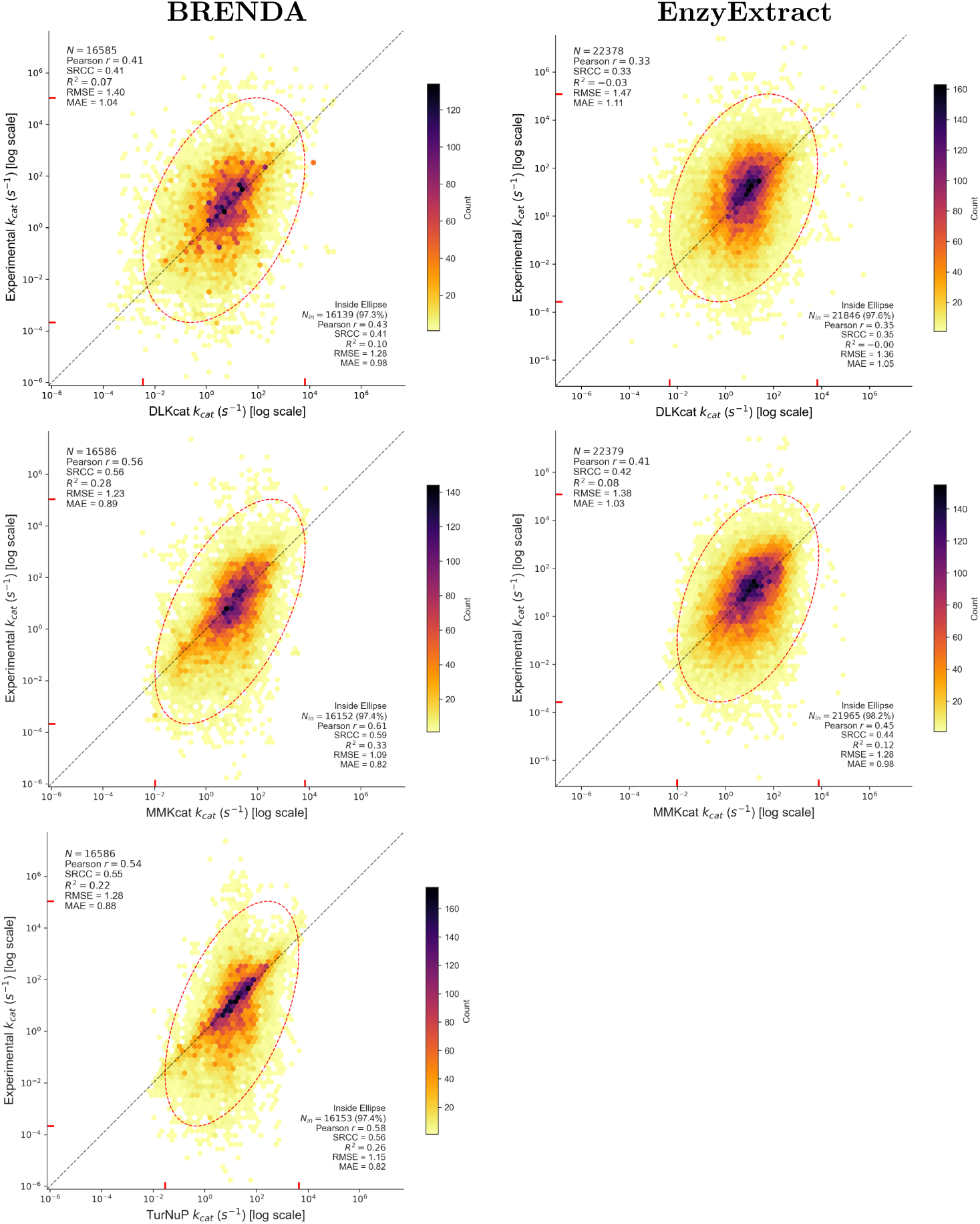
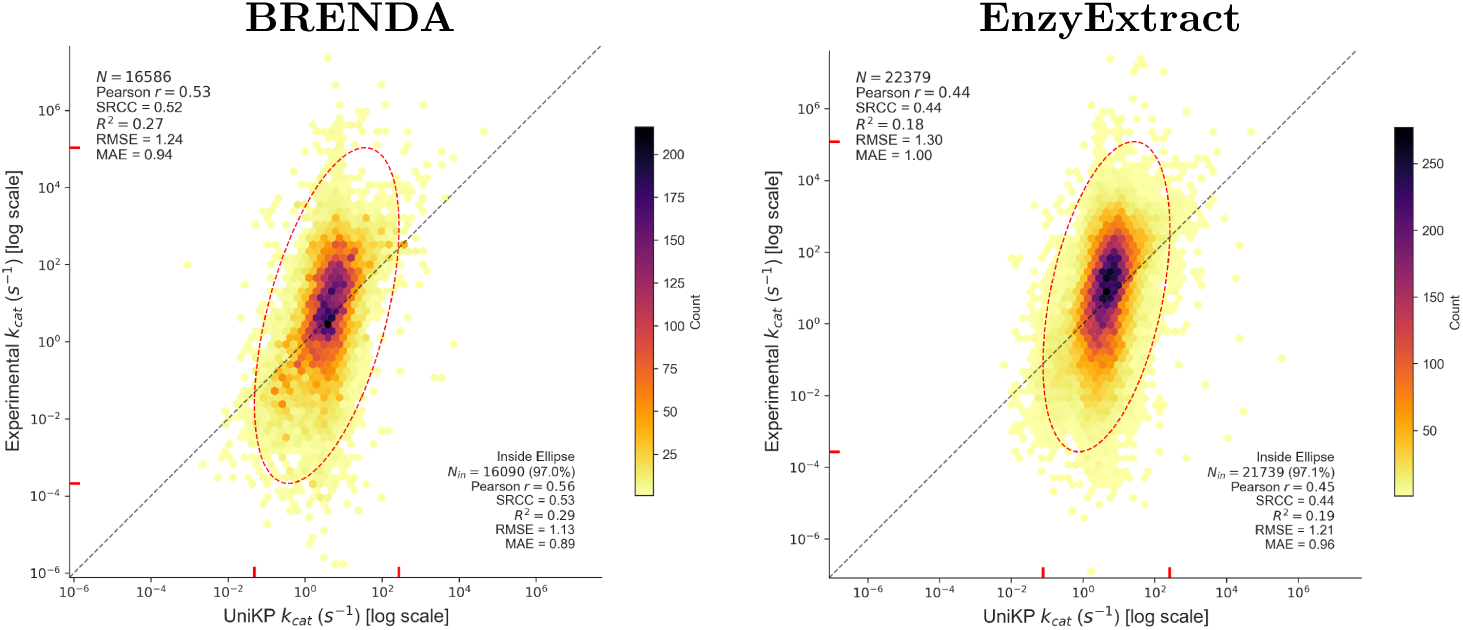
Predicted-versus-experimental k_cat_ values for all evaluated tools. Scatter plots show predicted k_cat_ values on the *x*-axis and experimentally measured k_cat_ values on the *y*-axis, both on a logarithmic scale. Rows correspond to prediction tools, ordered alphabetically, and columns correspond to the BRENDA-derived (left) and EnzyExtract (right) benchmarks. Point density is indicated by the color scale, and the diagonal dashed line denotes perfect agreement between predicted and experimental values. The overlaid ellipse represents a three-standard-deviation covariance ellipse of the predicted and experimental k_cat_ values. Red ticks on the *x*- and *y*-axes indicate the projections of the ellipse onto the respective axes. Performance statistics are reported for the complete dataset and for observations within the ellipse.

**Table SI 3:**
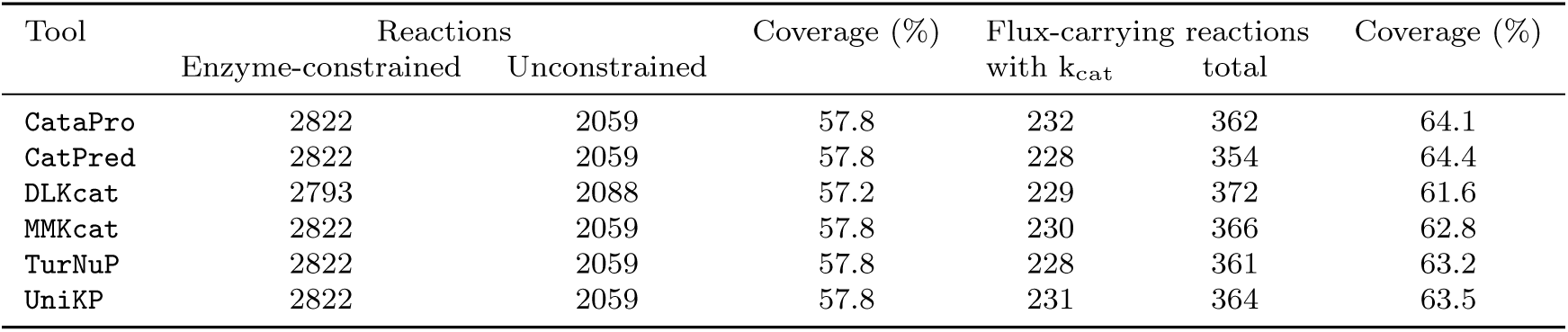
Coverage of k_cat_ parameterization in the tool-specific ecGEMs. The processed Yeast9 model contains 7122 irreversible reactions, of which 4881 are gene-associated. Reaction coverage is calculated relative to these gene-associated reactions. Flux-carrying reactions are defined as reactions with non-zero flux in the pFBA solution for glucose minimal medium, and flux-carrying coverage is calculated relative to the total number of flux-carrying reactions in each tool-specific ecGEM.

