## Supplementary Material for "Beyond benchmark accuracy: machine-learning turnover-number predictors require system-level validation"

### Supporting information

**Table SI 1**: Training-set sequence similarity of the BRENDA and EnzyExtract benchmarks. For each unique benchmark sequence, the closest sequence in the corresponding predictor training set was identified by global pairwise alignment. Values give the number of benchmark sequences within mutually exclusive best-match identity bins. Bin ranges are given in mathematical interval notation, with e.g. $[50, 70)\%$ indicating $50\,\% \leq x < 70\,\%$. Exact matches correspond to a best-match identity of 100 %. Mean identity gives the mean best-match sequence identity across all unique benchmark sequences (see Section 4 Materials and methods). In total, the BRENDA and EnzyExtract benchmarks contain 5854 and 9135 unique sequences, respectively. The two benchmark datasets themselves share 1892 exact sequences, corresponding to 20.71 % of unique EnzyExtract sequences and 32.32 % of unique BRENDA sequences.

| Tool | Benchmark | $[0, 50)\%$ | $[50, 70)\%$ | $[70, 90)\%$ | $[90, 100)\%$ | 100 % | Mean identity (%) |
|---|---|---|---|---|---|---|---|
| `CataPro` | BRENDA | 832 | 252 | 118 | 58 | 4594 | 89.44 |
| | EnzyExtract | 3975 | 1323 | 674 | 1016 | 2147 | 66.39 |
| `CatPred` | BRENDA | 859 | 291 | 129 | 129 | 4446 | 88.74 |
| | EnzyExtract | 3661 | 1266 | 742 | 1072 | 2394 | 68.57 |
| `DLKcat` / `UniKP`† | BRENDA | 2838 | 988 | 384 | 248 | 1396 | 62.87 |
| | EnzyExtract | 5220 | 1316 | 592 | 642 | 1365 | 58.48 |
| `MMKcat` | BRENDA | 2152 | 826 | 344 | 167 | 2365 | 70.61 |
| | EnzyExtract | 4721 | 1283 | 572 | 791 | 1768 | 61.17 |
| `TurNuP` | BRENDA | 3069 | 807 | 252 | 135 | 1591 | 62.16 |
| | EnzyExtract | 6302 | 1247 | 370 | 428 | 788 | 51.83 |

† The `UniKP` training set was inherited from `DLKcat`. After reduction to unique enzyme sequences for the present analysis, the two training sets were identical and therefore yielded the same sequence-overlap statistics.

**Table SI 2**: **Predicted-versus-experimental $k_{cat}$ values for all evaluated tools.** Scatter plots show predicted $k_{cat}$ values on the $x$-axis and experimentally measured $k_{cat}$ values on the $y$-axis, both on a logarithmic scale. Rows correspond to prediction tools, ordered alphabetically, and columns correspond to the BRENDA-derived (left) and EnzyExtract (right) benchmarks. Point density is indicated by the color scale, and the diagonal dashed line denotes perfect agreement between predicted and experimental values. The overlaid ellipse represents a three-standard-deviation covariance ellipse of the predicted and experimental $k_{cat}$ values. Red ticks on the $x$- and $y$-axes indicate the projections of the ellipse onto the respective axes. Performance statistics are reported for the complete dataset and for observations within the ellipse.

| **BRENDA** | **EnzyExtract** |
|---|---|

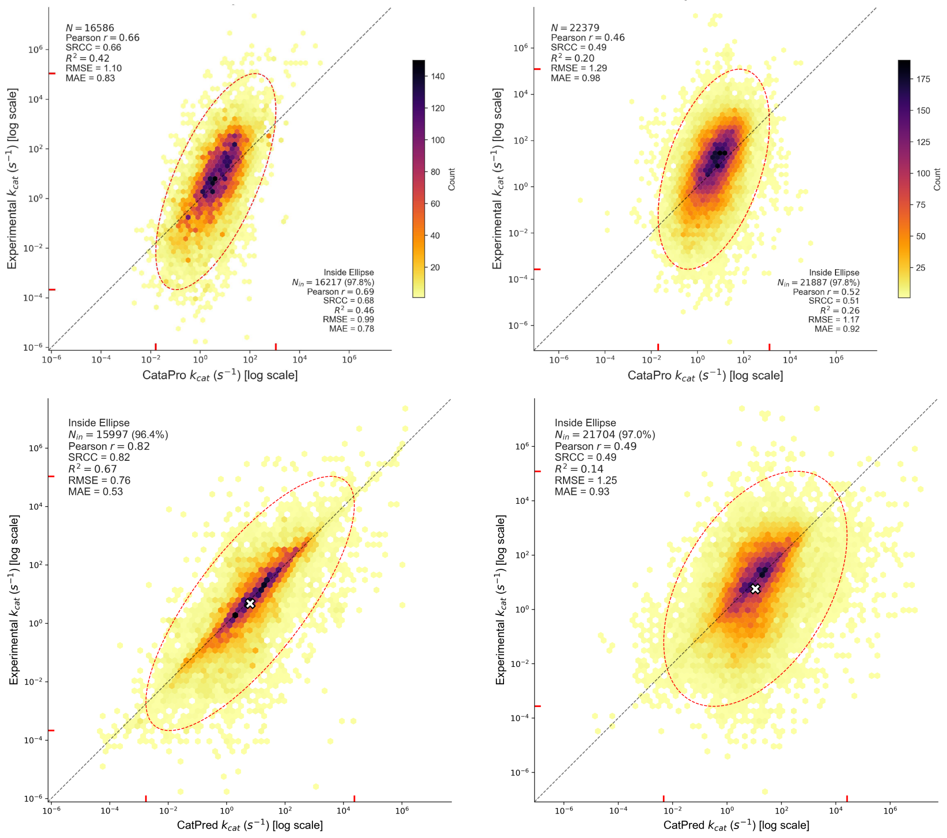


Continued on next page

**BRENDA** **EnzyExtract**

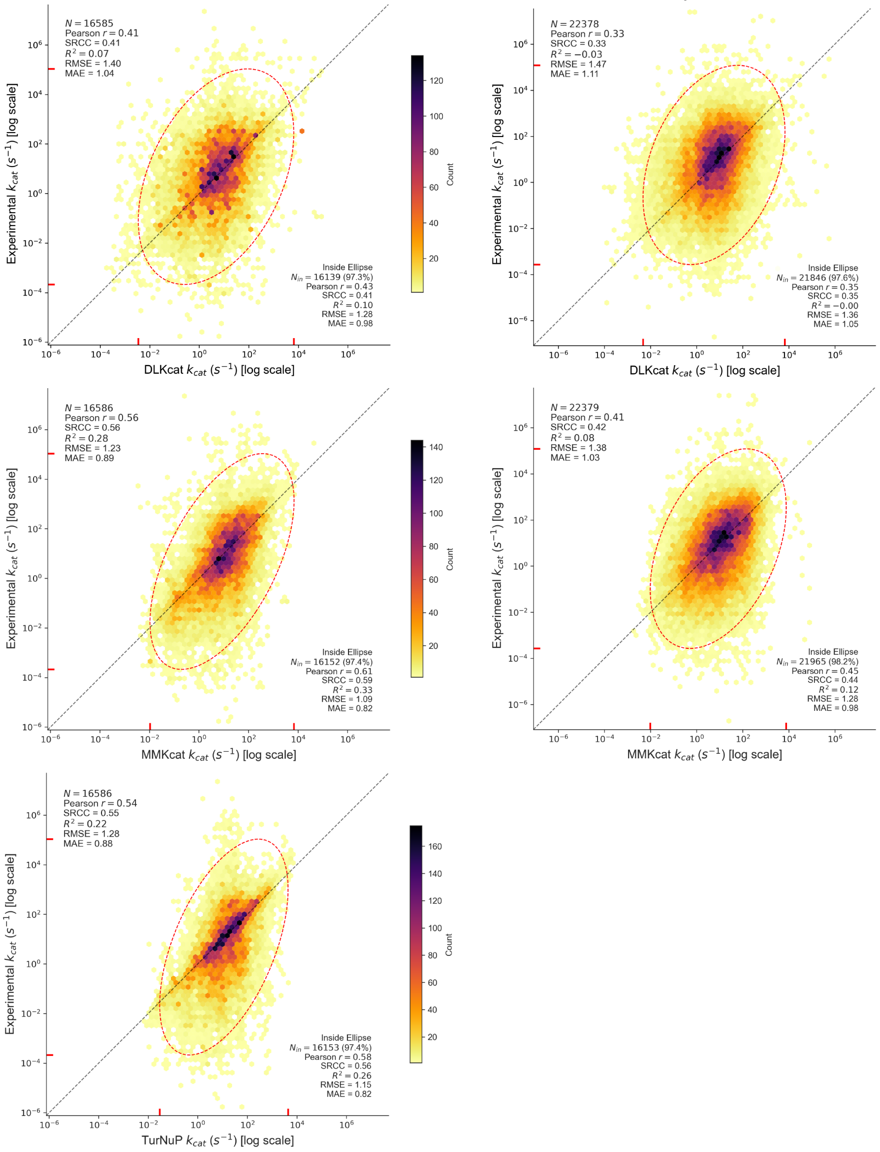


Continued on next page

Table SI 2 – continued from previous page

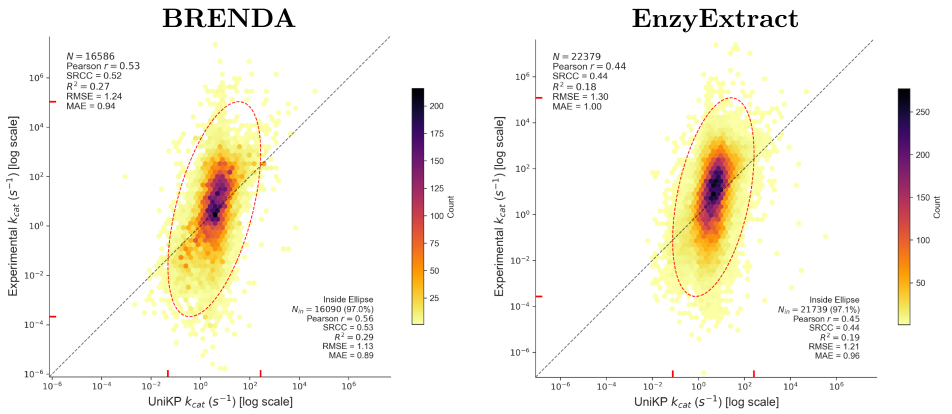


**Table SI 3**: Coverage of $k_{cat}$ parameterization in the tool-specific ecGEMs. The processed Yeast9 model contains 7122 irreversible reactions, of which 4881 are gene-associated. Reaction coverage is calculated relative to these gene-associated reactions. Flux-carrying reactions are defined as reactions with non-zero flux in the pFBA solution for glucose minimal medium, and flux-carrying coverage is calculated relative to the total number of flux-carrying reactions in each tool-specific ecGEM.

| Tool | Reactions | | Coverage (%) | Flux-carrying reactions | | Coverage (%) |
|---|---|---|---|---|---|---|
| | Enzyme-constrained | Unconstrained | | with $k_{cat}$ | total | |
| `CataPro` | 2822 | 2059 | 57.8 | 232 | 362 | 64.1 |
| `CatPred` | 2822 | 2059 | 57.8 | 228 | 354 | 64.4 |
| `DLKcat` | 2793 | 2088 | 57.2 | 229 | 372 | 61.6 |
| `MMKcat` | 2822 | 2059 | 57.8 | 230 | 366 | 62.8 |
| `TurNuP` | 2822 | 2059 | 57.8 | 228 | 361 | 63.2 |
| `UniKP` | 2822 | 2059 | 57.8 | 231 | 364 | 63.5 |

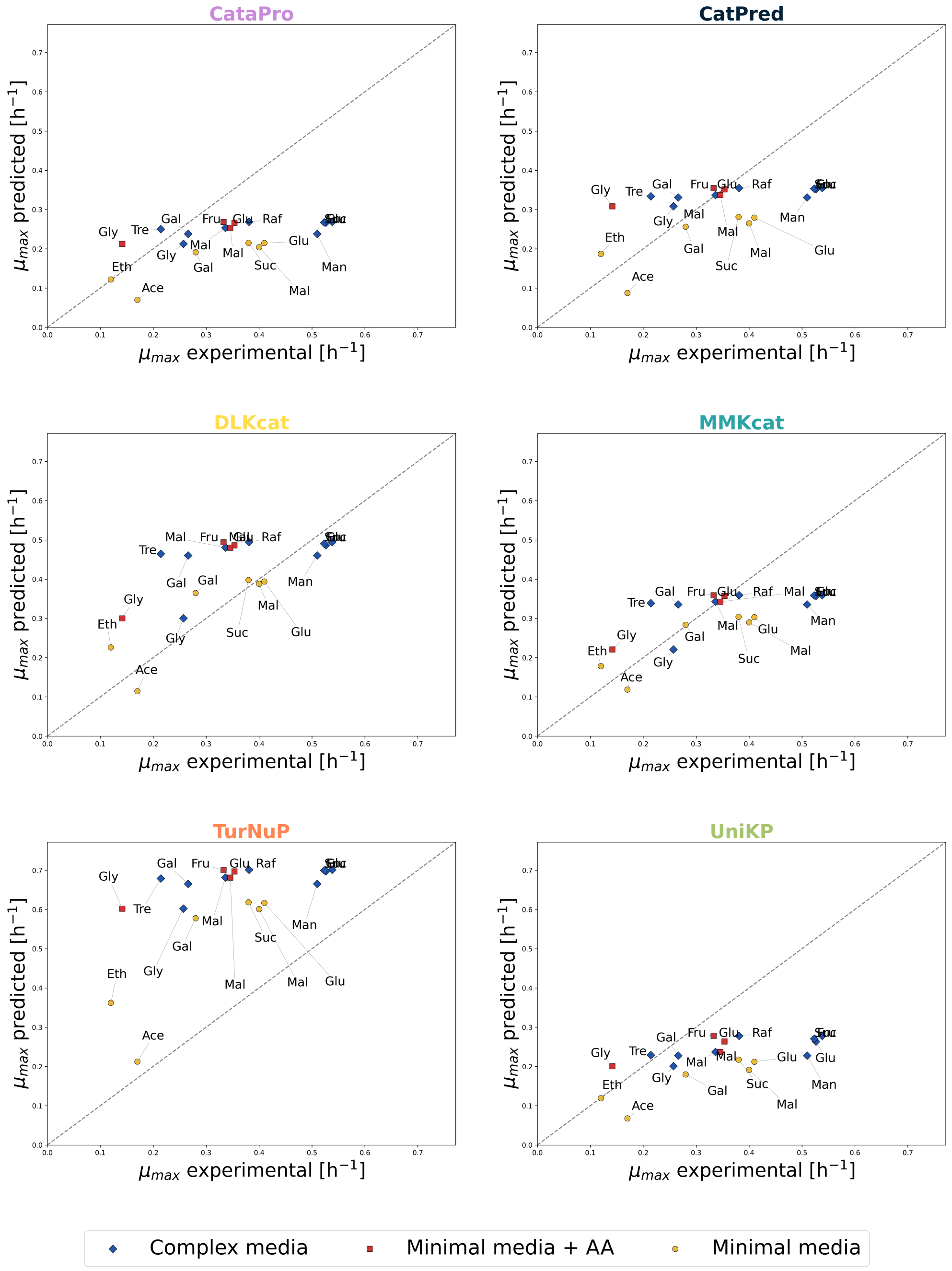


**Fig. SI 1**: Accuracy of ML-parameterized ecGEMs across multiple conditions. Predicted versus experimental maximum growth rates across carbon sources and medium compositions. Points represent individual conditions; marker color and shape indicate medium composition, and labels denote carbon source.
